# Empirical Analysis of Phylogenetic Quasi-Terraces

**DOI:** 10.1101/810309

**Authors:** Paula Breitling, Alexandros Stamatakis, Olga Chernomor, Ben Bettisworth, Lukasz Reszczynski

## Abstract

Terraces in phylogenetic tree space are, among other things, important for the design of tree space search strategies. While the phenomenon of phylogenetic terraces is already known for unlinked partition models on partitioned phylogenomic data sets, it has not yet been studied if an analogous structure is present under linked and scaled partition models. To this end, we analyze aspects such as the log-likelihood distributions, likelihood-based significance tests, and nearest neighborhood interchanges on the trees residing on a terrace and compare their distributions among unlinked, linked, and scaled partition models. Our study shows that there exists a terrace-like structure under linked and scaled partition models as well. We denote this phenomenon as quasi-terrace. Therefore quasi-terraces should be taken into account in the design of tree search algorithms as well as when reporting results on ‘the’ final tree topology in empirical phylogenetic studies.

## 1 Introduction

Phylogenetic trees are widely used to explain and visualize the evolutionary relationships of species. One approach to reconstructing phylogenies consists in using a supermatrix, which is a collection of multiple partitions, typically representing genes that are concatenated into one multiple sequence alignment. The alignment (matrix) size is determined by the number of homologous sites (columns) and the number of taxa (rows), often also denoted as species or sequences. Hence, a partition is a subset of alignment sites. A common problem with these large phylogenomic alignments are missing data. That is, a taxon can have no data present in a specific partition. This can either be due to errors in the sequencing process or because some species simply do not have data in this partition (do not have the specific gene) or if the gene has not been sampled yet (Kearney, 2002; Nyakatura & Bininda-Emonds, 2012; Wiens & Morrill, 2011). This type of missing data complicates phylogenetic tree inference under likelihood-based criteria (maximum likelihood (ML) or Bayesian inference) and under parsimony.

Two recent studies (Chernomor, von Haeseler, & Minh, 2016; Dobrin, Zwickl, & Sanderson, 2018) have analyzed a wide range of empirical data sets highlighting the importance and prevalence of terraces in tree in large contemporary phylogenomic studies. Dobrin et al. (2018) have primarily focused on the presence of terraces for published alignments and discussed their properties. Among others, for each data set, they provide the theoretical minimum number of genes that would be necessary to reduce the terrace sizes to just one tree. Moreover, the authors also discussed implications of terraces on bootstrap values.

The second study (Chernomor et al., 2016) introduced a production level implementation of techniques to account for terraces during tree searches in ML inference under the most general partition model, that is, the unlinked branch (UB) partition model (Z. Yang, 1996). The authors also introduced techniques for saving computational time in the presence of missing sequences under more restrictive linked (LB) and scaled branch (SB) partition models (Z. Yang, 1996). They obtained substantial speedups for all data sets tested under all three partition models. Awareness about quasi-terraces could help to further improve these results under the LB and SB models. During the search one can simply use one representative tree from a quasi-terrace to calculate its log-likelihood (LnL) score and treat it as an approximation of the log-likelihood scores for all other trees on the quasi-terrace. This will increase the speedup under LB and SB models by the same fraction as currently achieved under the UB partition model using terrace-aware tree inference procedures (in some cases up to 4.5 times faster, than in terrace-unaware computations).

## 2 Definitions

First we introduce some formal definitions and concepts. Let *T* and *Y* denote a tree and a partition, respectively. The induced partition tree, denoted by *T|Y*, is obtained from *T* by removing species, which have no data present in partition *Y* (Sanderson, McMahon, & Steel, 2011). A set of all trees that have an identical set of induced partition trees for all *k* partitions is denoted by *P* = {*T*_1_, *T*_2_,…, *T*_*N*_} (also called a *stand* in Sanderson, McMahon, Stamatakis, Zwickl, and Steel (2015)). This set is fully determined by the presence/absence matrix (presence/absence of genes per species/partition) and one representative tree *T*, which can be any of *T*_1_,…, *T*_*N*_.

For a given alignment (supermatrix) the tree space is partitioned into multiple disjoint *P* sets with (often) different sizes. The size (the number of trees in *P*) can vary between 1 tree (a trivial set) and more than a billion of trees (Sanderson et al., 2011). Note that, for a given set *P*, currently available algorithms can compute its size and enumerate all trees, only if the alignment contains a so-called comprehensive taxon (Biczok et al., 2018). That is, there is at least one taxon that has data for all partitions.

The three partition models (LB, SB, and UB; (Z. Yang, 1996)) differ in the way they optimize branch lengths. Under UB, a separate independent set of branch lengths is estimated for each partition in the phylogenomic data set. Under LB, a single branch length over all partitions is being estimated. Under SB, the same underlying branch lengths are scaled via a single parameter for each partition. Under all models, the LnL of a tree is computed as the sum of the LnLs over all induced partition trees. When the branch lengths are estimated independently for each induced partition tree, their LnLs, and as a result, also their sum are identical for all trees in *P*. That is, under the UB partition model, all trees in *P* have an identical LnL (Sanderson et al., 2015, 2011). In this case *P* is called a terrace. From theory we know, that under LB and SB, trees in *P* have distinct LnL (Sanderson et al., 2015). However, we expect that the LnL scores will not be significantly different. We call the set *P* under LB and SB a quasi-terrace.

As many phylogenetic studies of large empirical alignments use LB and SB partition models (Duchêne et al., 2018; Rosa, Melo, & Barbeitos, 2019; Wang, Susko, & Roger, 2019), the phenomenon of quasi-terraces is highly relevant for contemporary analyses. LB and SB partition models induce a substantially smaller number of free model parameters compared to UB. Consequently, they are less computationally expensive. Moreover, for the same reason, LB and SB are less prone to overfitting. For instance, it appears more reasonable to deploy the SB model to account for different evolutionary rates for partitions derived from the 1st, 2nd, and 3rd codon position, rather than using a generic UB partition model.

In this paper we conduct a study of 7 published empirical phylogenomic data sets to assess if quasi-terraces exist.

## 3 Data sets

Two recently published papers (Chernomor et al., 2016; Dobrin et al., 2018) addressed different aspects of terraces in tree space and collected a wide range of empirical alignments with missing data.

Dobrin et al. (2018) investigated how frequently terraces occur in empirical phylogenomic studies and analyzed data sets from 12 different published studies (Burleigh, Kimball, & Braun, 2015; Meredith et al., 2011; Miadlikowska et al., 2014; Misof et al., 2014; Rabosky, Donnellan, Grundler, & Lovette, 2014; Shi & Rabosky, 2015; Soltis et al., 2013; Springer et al., 2012; Tolley, Townsend, & Vences, 2013; Wickett et al., 2014; Y. Yang et al., 2015; Zanne et al., 2014). In total, they analyzed 23 DNA and 3 protein alignments. The terrace sizes among the data sets varied between a single tree and 1.30 *×* 10^388^ trees.

Chernomor et al. (2016) analyzed 12 empirical data sets (9 DNA and 3 protein alignments) published in (Bouchenak-Khelladi et al., 2008; Dell’Ampio et al., 2013; Fabre, Rodrigues, & Douzery, 2009; Hinchliff & Roalson, 2012; Nyakatura & Bininda-Emonds, 2012; Pyron et al., 2011; Springer et al., 2012; Stamatakis & Alachiotis, 2010; Van Der Linde, Houle, Spicer, & Steppan, 2010). The terraces corresponding to ML trees for the data sets from Chernomor et al. (2016) are either trivial (comprising just one tree) or extremely large (in some cases, exceeding 850, 000 trees).

In our study of quasi-terraces we use 7 phylogenomic DNA data sets from (Dobrin et al., 2018). Table 1 provides an overview of the analyzed data sets. The number of taxa varies from 106 to 213 and the number of partitions varies from 4 to 6. Other data sets from (Chernomor et al., 2016; Dobrin et al., 2018) had to be excluded, either because the terraces of interest were trivial (comprising just one tree), or excessively large (*>* 588, 735 trees) to be analyzed within the 24 hour job run time limit on the cluster.

**Table 1:** Overview of data sets

| Data set | Datatype | #Taxa | #Partitions | #Sites | Reference |
| --- | --- | --- | --- | --- | --- |
| Eucalyptus* | DNA | 136 | 4 | 6,205 | (Zanne et al., 2014) |
| Euphorbia* | DNA | 131 | 6 | 9,154 | (Zanne et al., 2014) |
| Primula* | DNA | 185 | 5 | 7,321 | (Zanne et al., 2014) |
| Rabosky.scincids | DNA | 213 | 6 | 5,373 | (Rabosky et al., 2014) |
| Ranunculus* | DNA | 170 | 6 | 10,799 | (Zanne et al., 2014) |
| Rhododendron* | DNA | 117 | 5 | 7,321 | (Zanne et al., 2014) |
| Szygium* | DNA | 106 | 4 | 5,815 | (Zanne et al., 2014) |

\*: *Subsampled data sets. Data sets were subsampled until they contained a comprehensive taxon. See Section 4.1 for details*

## 4 Experimental setup

For our quantitative analysis, we first designed a data preparation and analysis pipeline, so that every data set is prepared and analyzed in exactly the same way to obtain comparable results. We first outline the basic steps of our pipeline and subsequently discuss each of them in more detail.

For each data set we conducted the following steps (see also Table 2):

**Table 2:**
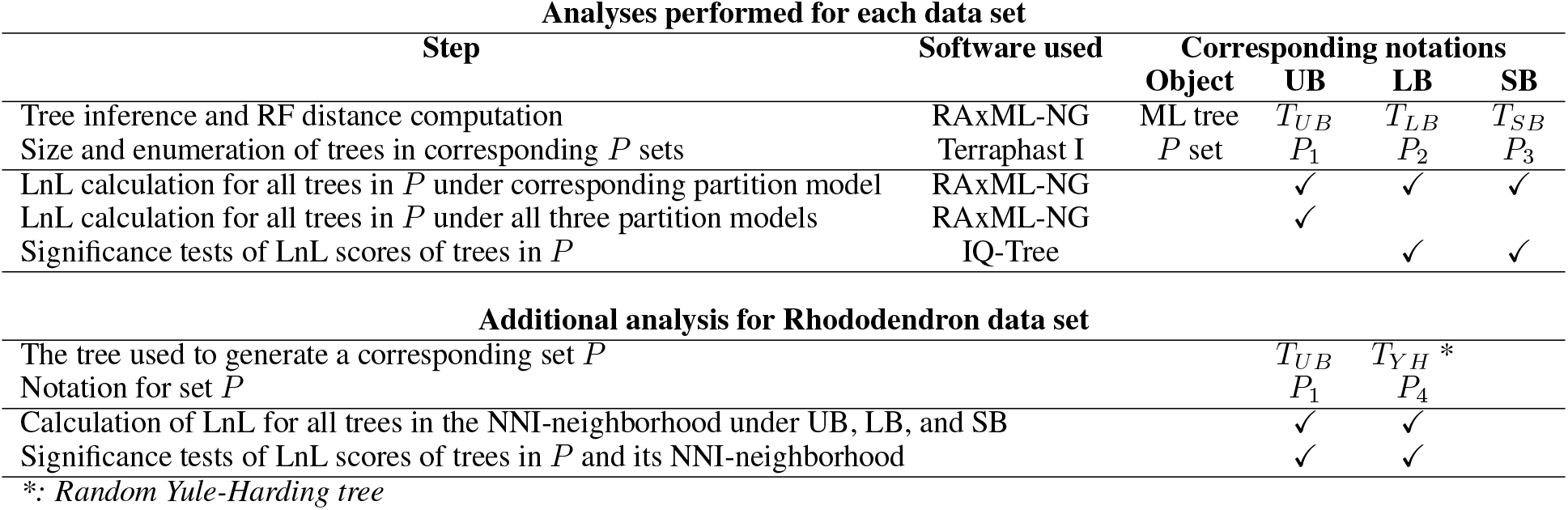
Analysis steps and notations

1. Data preprocessing: Create a PHYLIP formatted alignment, a binary presence/absence matrix (indicating which species has data for which partition), and a partition file (defining which columns/sites of the multiple sequence alignment belong to which partition) from the original NEXUS input files.
2. Conduct ML tree searches under UB, LB, and SB (inferred ML trees are denoted as *T*_*UB*_, *T*_*LB*_, and *T*_*SB*_, respectively).
3. For each ML tree from the previous step determine if the corresponding *P* set has size greater than 1 and enumerate the trees in *P* (for notations see Table 2).
4. Calculate the LnL scores of all trees in the respective *P* set under the corresponding partition model (*P*_1_ under UB, *P*_2_ under LB, *P*_3_ under SB).
5. Additionally, for *P*_1_ calculate the LnL score under the LB and the SB partition models.
6. Further analyses:

a. Calculate RF-distance (Robinson & Foulds, 1981) between the best ML trees: *T_LB_*, *T_SB_*, and *T*_*UB*_.
b. Compute significance tests based on the LnL scores for the trees in *P*_2_ and *P*_3_ under LB and SB.

The additional analysis of the tree neighborhood of the Rhododendron data set comprised the following steps:

1. For *P*_1_ generate its tree neighborhood by nearest-neighbor interchange (NNI^1^): for each tree from *P*_1_ collect all trees one NNI away from it.
2. Compute LnL for all trees in the NNI-neighborhood of *P*_1_ under the UB, LB, and SB partition models.
3. Compute significance tests of the trees in *P*_1_ and their NNI neighborhood for LnL scores under UB, LB, and SB.
4. The same NNI neighborhood analysis (generation of one NNI neighboring trees, LnL calculation, and significance tests) was also performed for the set *P*_4_ generated by a random Yule–Harding (YH) tree (Harding, 1971)).

### 4.1 Data preprocessing

As we had a collection of NEXUS data files, but required other formats for our analyses, we initially transformed our data accordingly. To analyze (quasi-)terraces we required a binary presence/absence matrix, which describes for which species we have data in which partitions.

An important aspect for the downstream analysis of (quasi-)terraces is that all data sets need to comprise a comprehensive taxon. However, not all of our data sets comprised such a comprehensive taxon *a priori*. Therefore, we reduced (subsampled) some data sets by removing partitions, such that the reduced data set comprised at least one comprehensive taxon. We conduct this reduction as follows: for each taxon we count in how many partitions it has data and chose the taxon that has the largest number of partitions *with* data. Once such a taxon has been selected, we remove all partitions from the data set for which this taxon does not have data. When such a reduction is applied, we need to propagate it to all subsequent files used within the analysis pipeline.

### 4.2 Tree inference

We inferred the best-known ML trees under the UB, LB, and SB models with RAxML-NG (Kozlov, Darriba, Flouri, Morel, & Stamatakis, 2019) (denoted as *T*_*UB*_, *T*_*LB*_ and *T*_*SB*_). For each data set and each partition model, we performed 20 independent tree searches, and recorded the tree with the best LnL score among these 20 independent runs. We used 10 random trees and 10 randomized stepwise addition order parsimony trees as starting trees.

### 4.3 Enumeration of trees in *P* sets

We used Terraphast I (Biczok et al., 2018) to enumerate all trees and computed the sizes of *P*_1_, *P*_2_, and *P*_3_ (collection of trees with identical induced partition trees), generated by the corresponding ML trees *T*_*UB*_, *T*_*LB*_, and *T*_*SB*_. Recall that, under UB, any set *P* is a terrace, while under LB and SB, any set *P* is a quasi-terrace. Apart from the ML tree file, Terraphast I requires the aforementioned binary presence/absence matrix as input. Terraphast I then outputs the number of trees in *P* (terrace/quasi-terrace) and enumerates all trees in *P* in Newick format.

### 4.4 Calculation of log-likelihood scores

We used RAxML-NG to compute the LnL scores of all tree topologies in the respective *P* sets. The input is a PHYLIP alignment file, the corresponding partition file, and the terrace tree file (trees in the corresponding *P* set). We calculated the LnLs of the trees in *P*_1_, *P*_2_, and *P*_3_ under the UB, LB, and SB partition models. For the sake of comparison, we also calculated the LnLs of the trees in *P*_1_ (generated by using *T*_*UB*_) under the LB and SB models. Recall that, any set *P* is defined by the topology of the representative tree. Therefore, the terrace *P*_1_ generated using *T*_*UB*_ is also a quasi-terrace under LB and SB.

### 4.5 Further analyses

To assess the similarity of the best ML trees inferred under each partition model and if partitioning ‘does matter’ (i.e., yields a distinct tree topology), we computed the Robinson-Foulds (RF) distances (Robinson & Foulds, 1981) between *T*_*UB*_, *T*_*LB*_, and *T*_*SB*_.

To confirm our expectation that the trees on a quasi-terrace are indeed not significantly different from each other with respect to the LnL score, we conducted standard significance tests with IQ-Tree (Chernomor et al., 2016; Nguyen, Schmidt, von Haeseler, & Minh, 2014) for the trees in *P*_2_ and *P*_3_ and the LnL scores computed under the LB and SB partition models, respectively. As input we used the partition file, the trees in the respective *P* set, and the PHYLIP alignment file.

IQ-Tree implements the following common LnL-based significance tests:

- Bootstrap proportions using the RELL method (Kishino, Miyata, & Hasegawa, 1990)
- Kishino-Hasegawa test (one sided and weighted) (Kishino & Hasegawa, 1989)
- Shimodaira-Hasegawa test (weighted and unweighted) (Shimodaira & Hasegawa, 1999)
- Expected likelihood weight (Strimmer & Rambaut, 2002)
- Approximately unbiased (AU) test by Shimodaira (Shimodaira, 2002)

To explore the trees surrounding a (quasi-)terrace, on one of our data sets (Rhododendron) we also performed a NNI neighborhood analysis. For each tree in *P*_1_ we applied all possible NNIs to obtain a collection of all one-NNI neighboring trees surrounding the (quasi-)terrace. We then compared the trees in *P*_1_ with the generated NNI tree set. We performed this comparison by evaluating LnL scores under all three partition models.

We identified the range between the maximum and minimum LnL of the trees in *P*_1_ (i.e., trees on the (quasi-)terrace). Then, we computed the fraction of NNI trees falling within this range and the fraction of NNI trees with a LnL higher than the maximum LnL of the trees in *P*_1_. We also conducted LnL score significance tests on an extended tree set, containing, the trees in *P*_1_ as well as the NNI trees.

With this analysis, we intended to assess how trees on a quasi-terrace differ from those in its NNI neighborhood. In other words, we test if all trees in *P* behave similarly, that is, if the majority of the trees on a quasi-terrace are significantly better or significantly worse than the surrounding trees. If all trees in *P* do behave similarly, it suffices to only use one representative tree from the quasi-terrace to save time and not evaluate other trees in *P*, that essentially behave similarly to that representative.

Since *P*_1_ was obtained using the best-known ML tree, the difference in LnLs between trees on the quasi-terrace and their NNI-neighbors might be small, as the tree search has converged on a tree with a ‘good’ ML score. Therefore, to assess, if the LnL differences are more pronounced for a quasi-terrace that was encountered during the early stages of the tree search, we also performed an analysis of the NNI neighborhood for a set *P*_4_ generated by a random Yule-Harding tree.

## 5 Results

### 5.1 Comparison of LnL scores for the trees from *P*_1_ under UB, LB and SB models

Table 3 summarizes the results for the corresponding *P*_1_ sets for all tested alignments. The size of *P*_1_ varies from 3 to 1, 863 trees. The average LnL (Avg. LnL) of the trees in *P*_1_ computed under UB is better (higher) than under SB and in turn under LB for all data sets. This is expected as the number of free model parameters decreases. The standard deviation (Std. Dev) of LnL scores is near zero for all data sets under UB and also for three data sets (Ranunculus, Rhododendron, and Szygium) under LB and SB. For Eucalyptus and Euphorbia, the Std. Dev for LB and SB is considerably larger (between 2.14 and 4.58). For Primula and Rabosky.scincids, it ranges between 1.21 and 1.78. The relative RF distance (Rel. RF) between the best ML trees *T*_*UB*_, *T*_*LB*_ and *T*_*SB*_ is smaller than 0.25 for 6 out of 7 data sets. For Eucalyptus we observe the highest value of 0.5.

**Table 3:** Overview of results for *P*_1_ sets

| Data set | Size of $P_1$ | UB | Avg. LnL LB | SB | UB | Std. Dev LB | SB | Rel. RF |
| --- | --- | --- | --- | --- | --- | --- | --- | --- |
| Euphorbia | 1,863 | -39,580.3 | -41,201.1 | -40,173.6 | 0.0009 | 4.5787 | 3.7331 | 0.15 |
| Primula | 1,125 | -37,034.1 | -38,491.0 | -38,084.1 | 0.0006 | 1.2102 | 1.2684 | 0.15 |
| Eucalyptus | 267 | -10,902.7 | -11,263.9 | -11,098.0 | 0.0120 | 3.2669 | 2.1400 | 0.50 |
| Rhododendron | 27 | -17,830.8 | -18,216.8 | -18,181.8 | 0.0005 | 0.1619 | 0.1644 | 0.25 |
| Ranunculus | 9 | -28,709.8 | -29,674.1 | -29,447.7 | 0.0009 | 0.0922 | 0.1452 | 0.18 |
| Szygium | 5 | -11,931.6 | -12,214.3 | -12,125.0 | 0.0003 | 0.1604 | 0.0155 | 0.18 |
| Rabosky.scincids | 3 | -125,550.6 | -128,649.7 | -127,973.0 | 0.0006 | 1.7823 | 1.6070 | 0.13 |

Figure 1 provides a comparison between the LnL scores of the trees in set *P*_1_ for the Rhododendron data set (plots for other data sets are available in the appendix, Figures A.1–A.6). The LnL was computed under all partition models (LB, SB, and UB). Here, the *x* axis represents the trees in *P*_1_ in the order they are enumerated by Terraphast I. Figure 1a shows the LnLs as computed under UB model, which should, in principle, all be exactly identical. The slight deviations are due to numerical round of error propagation. The maximal difference amounts to 0.0012 LnL units. Note that, for the LnL scores under the SB and LB models shown in Figure 1b and Figure 1c, the differences between the maximum and the minimum LnLs are also comparatively small (0.5558 and 0.4683 LnL units, respectively). The same holds for Ranunculus and Szygium, where the LnL differences under all partition models are near zero. For the remaining four data sets (Eucalyptus, Euphorbia, Primula, and Rabosky.scincids) the differences in LnL under SB and LB models are larger and vary between 3.4 and 22.4 LnL units. To verify the apparently structured LnL variations that are visible in the plots (presumably connected with the order by which the recursive function in Terraphast I enumerates trees on a terrace), we repeated the calculation of the LnL scores with IQ-Tree. The results resemble those of RAxML-NG (data not shown).

**Figure 1:**
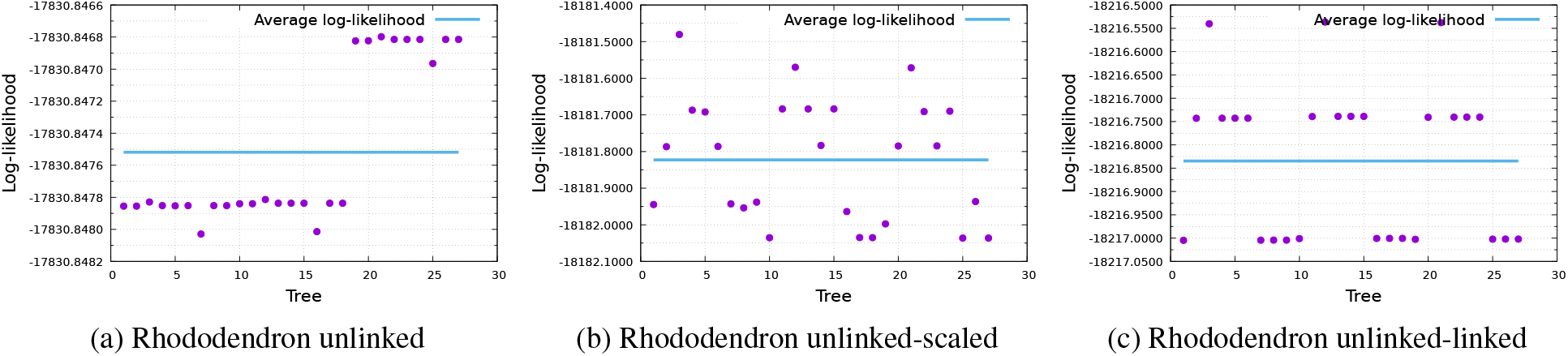
Log-likelihoods for Rhododendron data set under 1a UB, 1b SB, and 1c LB model

### 5.2 Significance tests for the trees from *P*_2_ and *P*_3_ with LnL under LB and SB partition models, respectively

Table 4 shows the results of the significance tests for the trees on quasi-terraces *P*_2_ and *P*_3_ and the LnL scores under LB and SB, respectively. For each data set, we report the fraction of trees on the quasi-terrace (*P*_2_ or *P*_3_), which are not significantly different with respect to their LnL from the best tree in the set (i.e., the tree with the highest LnL).

**Table 4:**
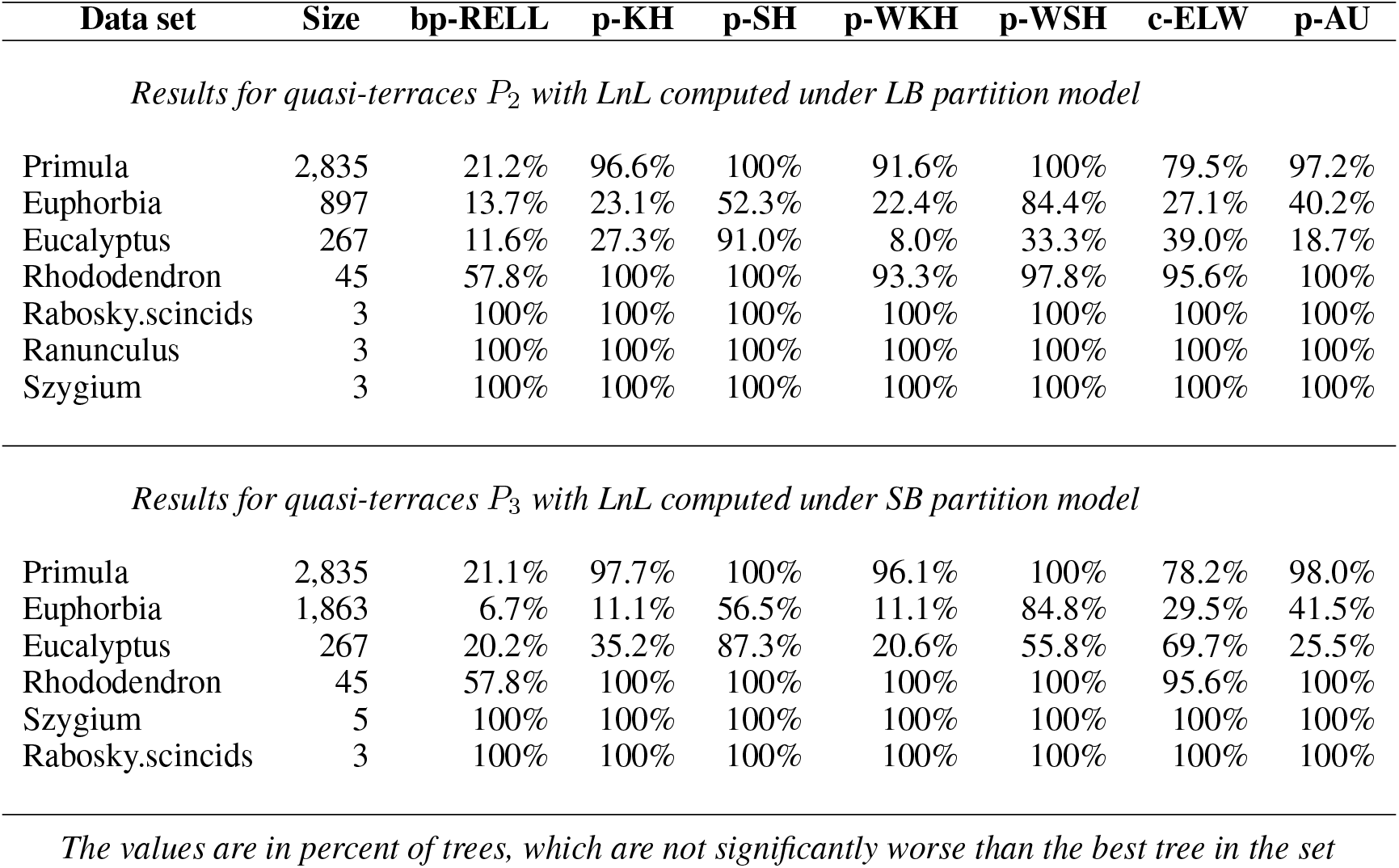
Significance tests for the trees on quasi-terraces

| Data set | Size | bp-RELL | p-KH | p-SH | p-WKH | p-WSH | c-ELW | p-AU |
| --- | --- | --- | --- | --- | --- | --- | --- | --- |
| *Results for quasi-terraces $P_2$ with LnL computed under LB partition model* |  |  |  |  |  |  |  |  |
| Primula | 2,835 | 21.2% | 96.6% | 100% | 91.6% | 100% | 79.5% | 97.2% |
| Euphorbia | 897 | 13.7% | 23.1% | 52.3% | 22.4% | 84.4% | 27.1% | 40.2% |
| Eucalyptus | 267 | 11.6% | 27.3% | 91.0% | 8.0% | 33.3% | 39.0% | 18.7% |
| Rhododendron | 45 | 57.8% | 100% | 100% | 93.3% | 97.8% | 95.6% | 100% |
| Rabosky.scincids | 3 | 100% | 100% | 100% | 100% | 100% | 100% | 100% |
| Ranunculus | 3 | 100% | 100% | 100% | 100% | 100% | 100% | 100% |
| Szygium | 3 | 100% | 100% | 100% | 100% | 100% | 100% | 100% |
| *Results for quasi-terraces $P_3$ with LnL computed under SB partition model* |  |  |  |  |  |  |  |  |
| Primula | 2,835 | 21.1% | 97.7% | 100% | 96.1% | 100% | 78.2% | 98.0% |
| Euphorbia | 1,863 | 6.7% | 11.1% | 56.5% | 11.1% | 84.8% | 29.5% | 41.5% |
| Eucalyptus | 267 | 20.2% | 35.2% | 87.3% | 20.6% | 55.8% | 69.7% | 25.5% |
| Rhododendron | 45 | 57.8% | 100% | 100% | 100% | 100% | 95.6% | 100% |
| Szygium | 5 | 100% | 100% | 100% | 100% | 100% | 100% | 100% |
| Rabosky.scincids | 3 | 100% | 100% | 100% | 100% | 100% | 100% | 100% |

*The values are in percent of trees, which are not significantly worse than the best tree in the set*

For the *P*_2_ sets and under the LB model, for 3 out of 7 data sets (Rabosky.scincids, Ranunculus, and Szygium) we find that all tests report that the trees are not significantly different from the best tree. For Primula and Rhododendron 6 tests suggest that the majority of the trees (79.5%-100%) are not significantly different from the best tree. For Eucalyptus and Euphorbia the results are inconclusive and the fractions of trees, which are not significantly different from the best tree, range from 8% to 91%.

The results for *P*_3_ quasi-terraces and LnL SCORES computed under SB resemble those of *P*_2_ and LB. Similarly, for Rabosky.scincids and Szygium all tests report that the trees in *P*_3_ are not significantly different from the best tree. For Primula and Rhododendron, 6 tests report that the majority of trees (78.2%-100%) are not significantly different from the best tree in the set. for Eucalyptus and Euphorbia the results are again inconclusive (the fractions of not significantly different trees vary from 6.7% to 84.8%). Note, that for the Ranunculus data set, *P*_3_ only contains one tree (i.e., the quasi-terrace is trivial). Therefore, significance tests are not available for Ranunculus.

### 5.3 NNI analysis for Rhododendron and a random Yule-Harding tree

For the NNI analysis we choose the Rhododendron data set. It contains sufficient number of trees in the *P* 1 set (27), but is, at the same time, still small enough in terms of taxa (117) such that the resulting NNI trees (in total 6141, with duplicates already removed) could be analyzed within the cluster runtime limit. For each partition model, LnL computations were completed within the runtime limits for a different number of NNI neighbors. Namely, 2389 NNI neighbors (38.9% out of total number of NNI trees) were evaluated under UB, 3720 NNI neighbors (60.58%) under LB, and 3705 (60.33%) under SB.

For all partition models, we calculated the fraction of NNI trees, whose LnL fall within the range between the maximum and minimum LnL of the trees *on* the (quasi-)terrace *P*_1_. These fractions amount to 15.8%, 30.2% and 27.9% under UB, LB, and SB, respectively. In addition, we also computed the fraction of NNI trees with a LnL greater than the maximum LnL for the trees in *P*_1_. The respective fractions of NNI trees are very small with only 3.9%, 1.4%, and 2.2% under UB, LB, and SB, respectively.

We performed the same analysis for a (quasi-)terrace generated using a random Yule-Harding tree. This random tree yields a set *P*_4_ of size 9 and the corresponding NNI neighborhood comprises 2048 trees (duplicates already removed). Here, we found that 18.9% NNI trees fall within the maximum-minimum LnL range of the trees *on* the (quasi-)terrace for UB, 30.3% for LB, and 51.5% for SB. These fractions for UB and LB models are similar to those obtained for *P*_1_. However, the difference is more pronounced under the SB model, where the fraction of NNI trees falling within the minimum-maximum LnL range of trees on the quasi-terrace increases from 27.9% to 51.5%. We also calculated NNI trees with LnL scores that are greater than those of the trees *on* the (quasi-)terrace *P*_4_. The results are: 38.2%, 36.5%, and 18.2% under UB, LB, and SB, respectively. These fractions are substantially greater than those for the (quasi-)terrace *P*_1_ generated using the ML tree. This is expected, as at toward the end of the tree search, the majority of trees surrounding the ML tree (and a (quasi-)terrace) are expected to have a LnL score that is similar to that of the ML tree.

For all three models we performed significance tests with IQ-Tree on the expanded set of trees including both, trees on the (quasi-)terrace, and trees in its neighborhood. As in the previous tests, we recorded the fraction of trees that are not significantly different from the best tree in the set. Table 5 summarizes the results: 27 trees in *P*_1_ are denoted in the Table 5 as “Terrace trees” and 6141 NNI trees as “NNI trees”. Under the UB model, 3 tests (bq-RELL, p-WKH, and p-WSH) report that all trees in *P*_1_ are significantly worse than the best tree in the set. The remaining 4 tests suggest the opposite (all trees in *P*_1_ are not significantly d ifferent). Similarly, under the LB and SB models, bq-RELL and p-WKH report that all trees in *P*_1_ are significantly worse than the best tree, while p-WSH reports that half of trees on quasi-terrace *P*_1_ is significantly worse than the best tree. Again, the remaining 4 tests suggest that all trees in *P*_1_ are not significantly different from the best tree. Overall, 6 tests suggested that the trees on the quasi-terrace behave alike. However, the trend (either all significantly or not significantly worse than the best tree) is not consistent among different tests. For the NNI trees, the results are inconclusive and the fraction of NNI trees that are not significantly worse than the best tree in the set varies substantially for all three branch length models.

**Table 5:** Significance tests for the trees on (quasi-)terraces and their NNI neighborhoods (Rhododendron data set)

| Model | Trees | bp-RELL | p-KH | p-SH | p-WKH | p-WSH | c-ELW | p-AU |
| --- | --- | --- | --- | --- | --- | --- | --- | --- |
| *Results for $P_1$ and its NNI neighborhood* |  |  |  |  |  |  |  |  |
| **UB** | All trees | 6.89% | 76.46% | 97.37% | 59.34% | 81.71% | 59.57% | 80.89% |
|  | Terrace trees | 0.00% | 100.00% | 100.00% | 0.00% | 0.00% | 100.00% | 100.00% |
|  | NNI trees | 6.92% | 76.36% | 97.36% | 59.60% | 82.07% | 59.39% | 80.80% |
| **LB** | All trees | 6.29% | 78.75% | 96.50% | 48.98% | 87.01% | 60.30% | 83.04% |
|  | Terrace trees | 0.00% | 100.00% | 100.00% | 0.00% | 55.56% | 100.00% | 100.00% |
|  | NNI trees | 6.32% | 78.65% | 96.48% | 49.19% | 87.15% | 61.12% | 82.97% |
| **SB** | All trees | 6.91% | 79.12% | 96.50% | 40.24% | 84.14% | 61.69% | 83.80% |
|  | Terrace trees | 0.00% | 100.00% | 100.00% | 0.00% | 74.07% | 100.00% | 100.00% |
|  | NNI trees | 6.94% | 79.03% | 96.48% | 40.42% | 84.19% | 61.52% | 83.73% |
| *Results for $P_4$ and its NNI neighborhood* |  |  |  |  |  |  |  |  |
| **UB** | All trees | 1.94% | 4.23% | 19.54% | 4.38% | 40.25% | 2.43% | 7.00% |
|  | Terrace trees | 0.00% | 0.00% | 0.00% | 0.00% | 0.00% | 0.00% | 0.00% |
|  | NNI trees | 1.95% | 4.25% | 19.63% | 4.39% | 40.43% | 2.44% | 7.03% |
| **UB-LB** | All trees | 1.56% | 3.89% | 13.08% | 2.04% | 37.92% | 1.99% | 2.92% |
|  | Terrace trees | 0.00% | 0.00% | 0.00% | 0.00% | 22.22% | 0.00% | 0.00% |
|  | NNI trees | 1.56% | 3.91% | 13.13% | 2.05% | 37.99% | 2.00% | 2.93% |
| **UB-SB** | All trees | 1.51% | 3.60% | 14.05% | 2.14% | 47.59% | 1.94% | 2.92% |
|  | Terrace trees | 0.00% | 0.00% | 0.00% | 0.00% | 55.56% | 0.00% | 0.00% |
|  | NNI trees | 1.51% | 3.61% | 14.11% | 2.15% | 47.56% | 1.95% | 2.93% |

*The values are in percent of trees, which are not significantly worse than the best tree in the set*

We also performed significance tests for the (quasi-)terrace *P*_4_ generated by a random tree and its NNI neighborhood (Table 5). There are 9 terrace trees, 2048 NNI trees, and hence a total of 2059 analyzed trees. Here, the behavior of the trees on the (quasi-) terraces is more consistent across different models and different tests. In contrast to the results for *P*_1_, under all three models 6 tests report that all trees in *P*_4_ are not significantly worse than the tree with the best LnL. The p-WSH test reported the same for the UB model, but also suggests that approximately half the trees from *P*_4_ are significantly worse than the best tree under the LB and SB models.

## 6 Discussion

In this study we analyzed 7 empirical data sets and their (quasi-)terraces as generated on the best-found ML trees. We compared the LnL scores of the trees on the (quasi-)terraces under all three partition models. The significance tests showed that trees on quasi-terraces are not significantly different from the best tree in the set with respect to their LnL scores. Therefore, during the tree search we can approximate the LnL scores of all trees on a quasi-terrace by evaluating the LnL of only one representative tree on that quasi-terrace.

For one of our data sets (Rhododendron), we also performed an analysis of the NNI neighborhood of the quasi-terraces *P*_1_ and *P*_4_, generated by the ML tree under the UB model (*T*_*UB*_) and by a random Yule-Harding tree, respectively. Significance tests on theses extended tree sets, including trees from the quasi-terrace *and* its NNI neighborhood, suggest that trees on a quasi-terrace behave similarly. This again implies that we can potentially use only one representative tree from the quasi-terrace during the tree search to accelerate the exploration of the tree space region containing that quasi-terrace. Such an approximation (using only one representative tree) can be beneficial, in particular during the initial phases of the tree search, to rapidly navigate out of regions with low LnL scores.

Our study constitutes a preliminary inspection of quasi-terraces and a more thorough study is required to confirm our results at a broader scale. Nevertheless, our findings suggest that quasi-terraces indeed exist and can be helpful for increasing the efficiency of tree search algorithms.

## Footnotes

1 Nearest-neighbor interchange is the simplest topological rearrangement operation, when one tree is obtained from another by changing subtrees across an internal branch of the tree.

---

**Figure A.1:**
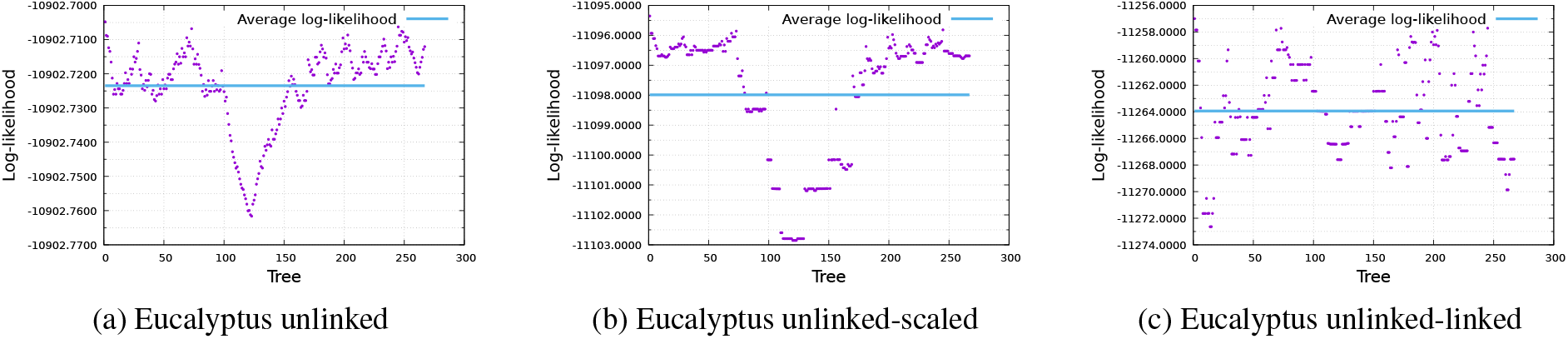
Log-likelihoods for the data set Eucalyptus under A.1a UB, A.1b UB-SB, and A.1c UB-LB model

**Figure A.2:**
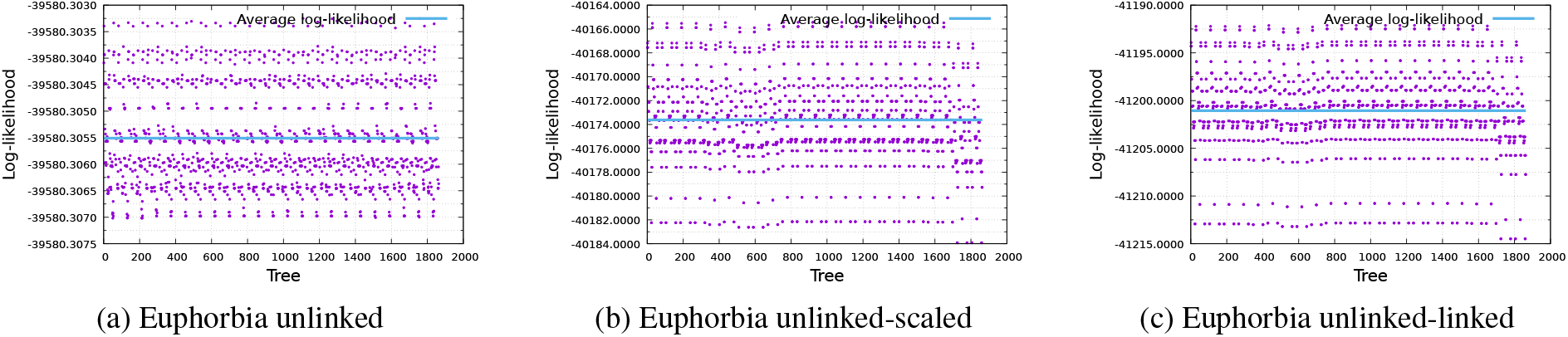
Log-likelihoods for the data set Euphorbia under A.2a UB, A.2b UB-SB, and A.2c UB-LB model

**Figure A.3:**
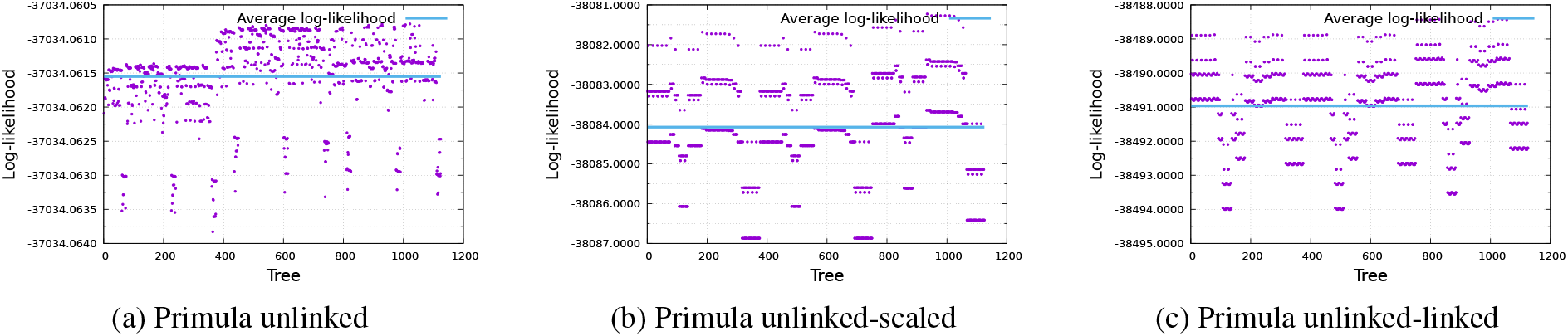
Log-likelihoods for the data set Primula under A.3a UB, A.3b UB-SB, and A.3c UB-LB model

**Figure A.4:**
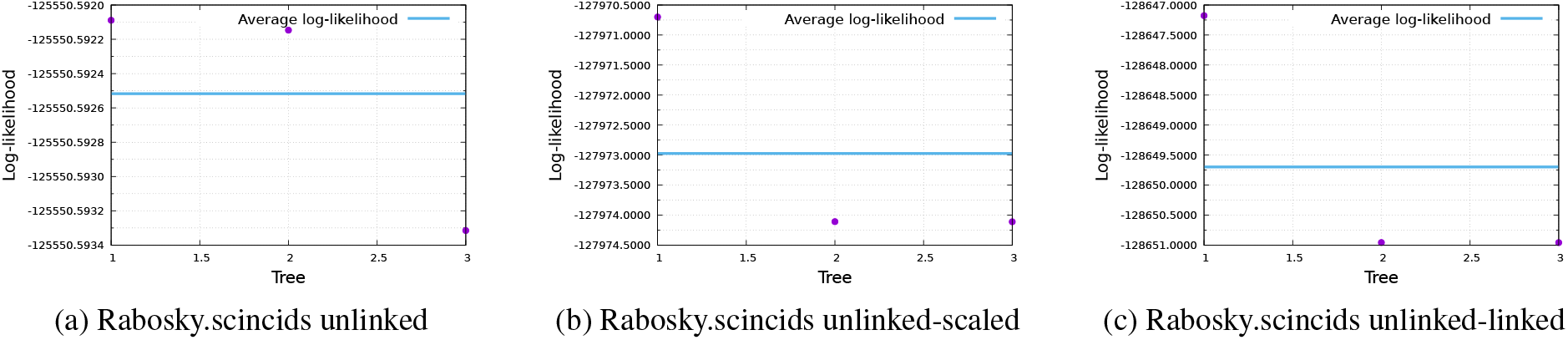
Log-likelihoods for the data set Rabosky.scincids under A.4a UB, A.4b UB-SB, and A.4c UB-LB model

**Figure A.5:**
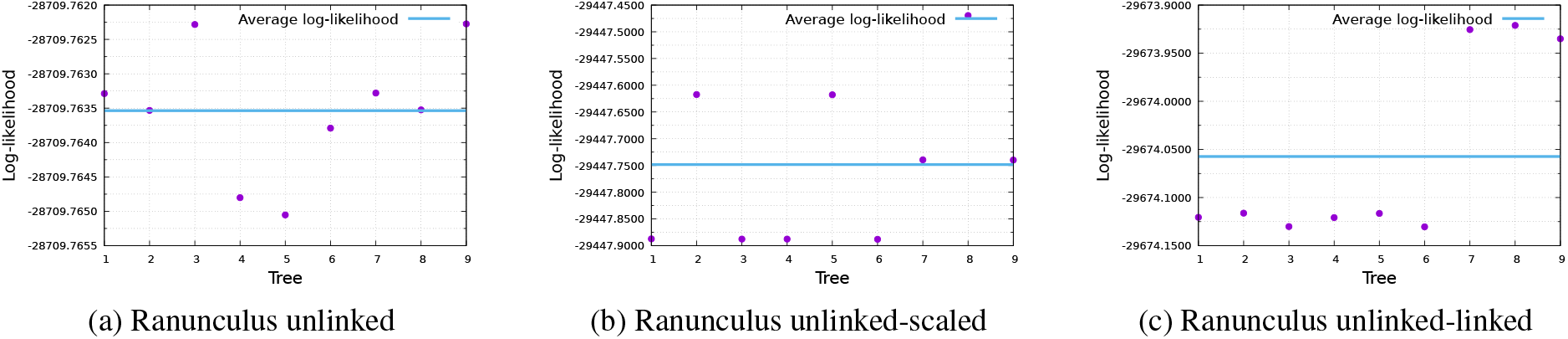
Log-likelihoods for the data set Ranunculus under A.5a UB, A.5b UB-SB, and A.5c UB-LB model

**Figure A.6:**
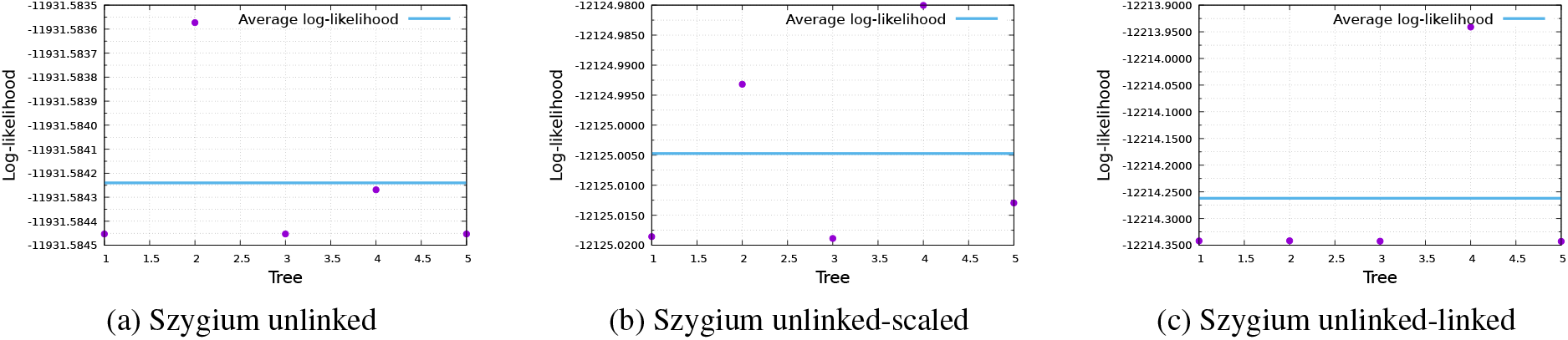
Log-likelihoods for the data set Szygium under A.6a UB, A.6b UB-SB, and A.6c UB-LB model

